# Differential Expression Analysis of miRNAs and mRNAs in Epilepsy Uncovers Potential Biomarkers

**DOI:** 10.1101/2023.09.11.557132

**Authors:** Fatma El Abed, Ghada Baraket, Marion N. Nyamari, Careen Naitore, Olaitan I. Awe

**Affiliations:** Department of Biology, Faculty of Sciences, University of Manar, Tunis, Tunisia; Department of Biological Sciences, Pwani University, Kilifi, Kenya; Jomo Kenyatta University of Agriculture and Technology, Nairobi, Kenya; Department of Computer Science, University of Ibadan, Ibadan, Oyo State, Nigeria; African Society for Bioinformatics and Computational Biology, Cape Town, South Africa

**Keywords:** Epilepsy, Seizure, miRNA, mRNA, Differentially Expressed Genes

## Abstract

Epilepsy is a neurological disease defined by episodes of synchronous convulsions. Recently, miRNAs have been proven as promising biomarkers for multiple ailments like tumors and neurodegenerative disorders; their role in epilepsy is still unclear. This study aimed to understand the involvement of miRNAs in the disease and to detect the potential biomarkers for the treatment of epilepsy.

RNA transcripts, and miRNA from brain tissue and plasma small extracellular vesicle samples of epileptogenic patients from 6 different studies downloaded from the NCBI sequence read archive (SRA) were analyzed with particular interest in genes that might be involved in epilepsy. Alignment of transcripts to hg38 was done using HISAT2 and the raw counts were generated using HTseq-count. miRNA genes were identified using miRDeep2. EdgeR and GEO2 were used to identify DEGs for both mRNA and miRNA datasets. Finally, TargetScan web tool was used to predict potentially significantly expressed mRNA target genes using the identified miRNA genes.

Analysis of these datasets revealed target genes in epilepsy and their associated miRNAs. SIX4 and KCTD7 were under-expressed in epileptogenic zones of the brain compared to the irritative zone. CABP1, SLC20A1 and SLC35G1 were under-expressed in brain tissues. Hsa-miR-27a-3p was identified as a regulator of CABP1 expression, hsa-let-7b-5p regulates SLC20A1 while hsa-miR-15a-5p and hsa-miR-195-5p are regulators for SLC35G1. These observations highlight the importance of miRNAs as novel biomarkers of epilepsy.

Understanding and controlling these regulatory interactions may help to define potential therapies for epilepsy. This would also help to better understand miRNA-mediated gene regulation in epilepsy.

## Introduction

Epilepsy is a complex disease that results from various abnormalities in the brain function caused by extraordinarily synchronized neural cell discharge. Clinically, it’s expressed as recurrent convulsions which may cause a loss of consciousness and can seriously affect daily activity of people with epilepsy [1]. Seizures are the result of an imbalance between excitatory and inhibitory neurotransmission. It can be focal seizure affecting a limited area or cerebral hemisphere or it can be a generalized seizure which is bilateral [2]. There are about 70 million cases of epilepsy worldwide and 65% have an unknown etiology [3]. According to statistics, children can develop epilepsy at a rate of between thirty-three and eighty-two per 100,000 children annually [4]. Epilepsy does not affect all ages equally, its distribution is bimodal, with a first peak at 5-9 years and the second peak at about 80 years. There is no difference in the prevalence of epilepsy according to gender [5]. The current classification of epilepsy was created in 2017. It is a multilevel classification that is based on the wide variation of resources in the world and the diagnosis must include three levels as well as the etiology of epilepsy [6].

The classification of epilepsy begins with determining the type of seizure [6]. This step requires a clinician to define that it is an epileptic seizure and to distinguish it from another non-epileptic event. The convulsions can be “focal seizures”, “generalized seizures” and “seizures of unknown onset” [6]. The second step is to determine the type of disease. The type of epilepsy consists of two already known types (focal and generalized epilepsy) with a new category that is a combination of the two known types [6]. There is also an undetermined or unknown category [6]. The other categorization is based on the determination of the Epileptic Syndrome [6]. The epileptic syndrome is defined by a constant and non fortuitous association of clinical and paraclinical characteristics such as the type of seizures, the EEG and the imaging [6].

It also includes seizure-promoting factors, their variable occurrence according to the sleep-wake cycle and aspects that rely on age, such as the onset-age and remission [6].

Actually, the diagnosis is made by looking at the symptoms, the patient’s medical history, and electroencephalograms, neuroimaging, and ultrasonography but it is still not sufficient in some cases. That is why a genetic biomarker would have the potential to revolutionize identification and facilitate the appropriate therapy [7], [8], [9].

An epileptic seizure is initiated when a network of brain neurons is activated and becomes hyperexcitable, which means that the nerve cells responsible for conducting the nerve impulse are activated in a sustained manner and will generate simultaneous electrical activity described as hyper-synchronous [5].

In fact, the neuron is a cell that constitutes the functional unit of the nervous system. It ensures the transmission of electrical signals called nerve impulses. The membrane of the neuron receives excitatory synaptic signals (depolarizing) and inhibitory synaptic signals (hyperpolarizing). The potential of the neuronal membrane reaches a threshold value if the sum of the excitatory synaptic inputs exceeds that of the inhibitory inputs, the neuron will then discharge an action potential which corresponds to the opening of the ion channels thus allowing the entry of sodium [10]. For this, three major hypotheses have been proposed: either the disease results from a defect in inhibition or hyperexcitation or a modification of cellular properties such as channelopathy or channel disease [10].

The two main neurotransmitters involved in epilepsy are glutamate and GABA; a central nervous system’s main inhibitory neurotransmitter [11]. This molecule exerts its function according to specific receptors: GABA-A, GABA-B, GABA-C. The fixation of GABA on its receptor GABA-A activates an ionic channel allowing the entry of chlorine ions into the neuron, thus inducing hyperpolarization which produces an inhibitory signal to the cell and decreases the probability of an action potential [11]. When the GABA-B receptor is active, it activates a metabotropic receptor that is permeable to potassium ions and determines a slower inhibitory response [11]. All of these receptors have ion channels that open upon binding of their ligands allowing the entry of ions of sodium and calcium thus inducing an excitatory signal [12]. The activation (glutamate) and inhibition (GABA) of neurotransmitter systems being out of balance can lead to neuronal hyper-excitability and thus the onset of an epileptic seizure [12].

Small non-coding RNAs called miRNAs control the expression of genes by the degradation of mRNA or the inhibition of protein translation [13]. The interaction between miRNA and target mRNA is characterized by imperfect base pairing between the two RNA sequences. The strands of the miRNA often form bulges [14]. The sequence specificity, which allows the miRNA to know its target mRNA, is determined by 2 to 8 nucleotides of its 5’ region, called the seed sequence [14].

MicroRNAs have now been suggested for clinical diagnosis for a variety of disorders due to their serum stability, affordability, speed, and non-invasive qualities [15]. In the future, genomic medicine would rely largely on our ability to use sequencing data for clinical applications like biomarker discovery, viral pathogen evolution [16], SARS-CoV-2 variants classification [17], for other applications like screening for disorders in newborns [18], HIV-1 evolution in sub-Saharan Africa [78], prostate cancer biomarker discovery [79], malaria/CoVID-19 biomarker discovery [80], Chromatin Immunoprecipitation [81], and viral comparative genomics [82].

Although some authors have reported micro-RNAs that are connected to epilepsy, the majority of them have to do with glial regeneration, neuronal death, and the inflammation processes [6], [7], [4], [19], [20], [21].

The study aimed to i) determine differential expression of genes and miRNAs between disease and controls using the developed tools and ii) identify new miRNA linked to epilepsy and targeting new genes implicated in the onset of seizures.

## Methods

### 1. Datasets

In this study, we used publicly available NGS datasets available on the NCBI GEO database (https://www.ncbi.nlm.nih.gov/geo/) [28]. To obtain the data sets, we used a predetermined search strategy using keywords or terms of the medical subject headings such as “epilepsy” AND “cerebral” AND “cerebral tissues’’ AND “seizure” and the species filter was fixed to “homosapiens’’. In the next step, we first set the study type to “expression profiling data’’ in order to obtain mRNA expression data, and subsequently set it to “non-coding RNA profiling by array” and “non-coding RNA profiling by high throughput sequencing” for the miRNA data retrieval. Datasets obtained and used had the following GEO and SRA accession numbers; 1. mRNA data: GSE127871 (brain tissue samples), SRA study: <u>SRP187565</u>; GSE57585, GSE31718 (brain tissue samples), 2. miRNA data: GSE97365 (brain tissue samples), GSE193842 (plasma small extracellular vesicles), SRA study: SRP355454 3. miRNA + mRNA data: GSE 205661 (brain tissue samples).

The obtained datasets contain different information about patients and controls such as etiology, onset of seizures, frequency, age and sex which allows the user to create their own groups for the data.

Most of the data was generated from cerebral tissues which allowed us to compare the differential expression in neurons which are the original zone of epilepsy (Table 1).

**Table 1:** Summary of the datasets integrated for the prediction of gene targets.

| Datasets | Platform | Number of Samples | Number of Epilepsy Patients | Tissue type | Number of Controls | Country |
| --- | --- | --- | --- | --- | --- | --- |
| GSE127871 | GPL16791 | 12 | 12 | Brain tissue | 0 | USA |
| GSE 205661 | GPL18402<br>GPL19072 | 12 | 6 | Brain tissue | 6 | China |
| GSE 57585 | GPL6480 | 14 | 7 | Brain tissue | 7 | Brazil |
| GSE 193842 | GPL20301 | 33 | 23 | Plasma sEvs | 10 | China |
| GSE 31718 | GPL6254 | 10 | 10 | Brain tissue | 0 | China |
| GSE 97365 | GPL8786 | 10 | 5 | Brain tissue | 5 | Brazil |

### 2. Data Analysis

#### 2.1. Analysis of mRNA data (GSE127871)

Data was downloaded from the GEO database. Subsequently, using fastQC version 0.11.3., the sequencing data’s quality was evaluated [22] (Fig. 2).

Next, we used Cutadapt to eliminate poor quality reads as well as adaptor sequences following the threshold parameters described by Martin [48]. The validated mRNA reads were then aligned to the human reference genome (hg38) using Hisat2 [38]. Subsequently, we verified the quality of sequence alignments using samtools stats [42]. The mRNA SAM file generated from the aligned reads was converted to a BAM file for further processing using Samtools v1.3.1. [42]. To quantify the number of mRNA reads, we used the HTSeq-count package [56] to determine the copy number associated with each human gene. Subsequently we analyzed correlations between samples using the volcano plot function in EdgeR [60].

#### 2.2. Analysis of microRNA data (GSE193842)

The data was downloaded from the GEO database. Then, using fastQC version 0.11.3., the sequencing data’s quality was evaluated [22] and adapter sequences and poor quality sequences were removed by the Trimmomatic package [27]. Alignment of high quality reads was done using the Bowtie package [41] and we identified miRNAs using miRDeep2.pl [32]. To quantify the miRNAs, we used a quantifier.pl script available in miRDeep2 [32] (Fig 1).

**Fig 1.**
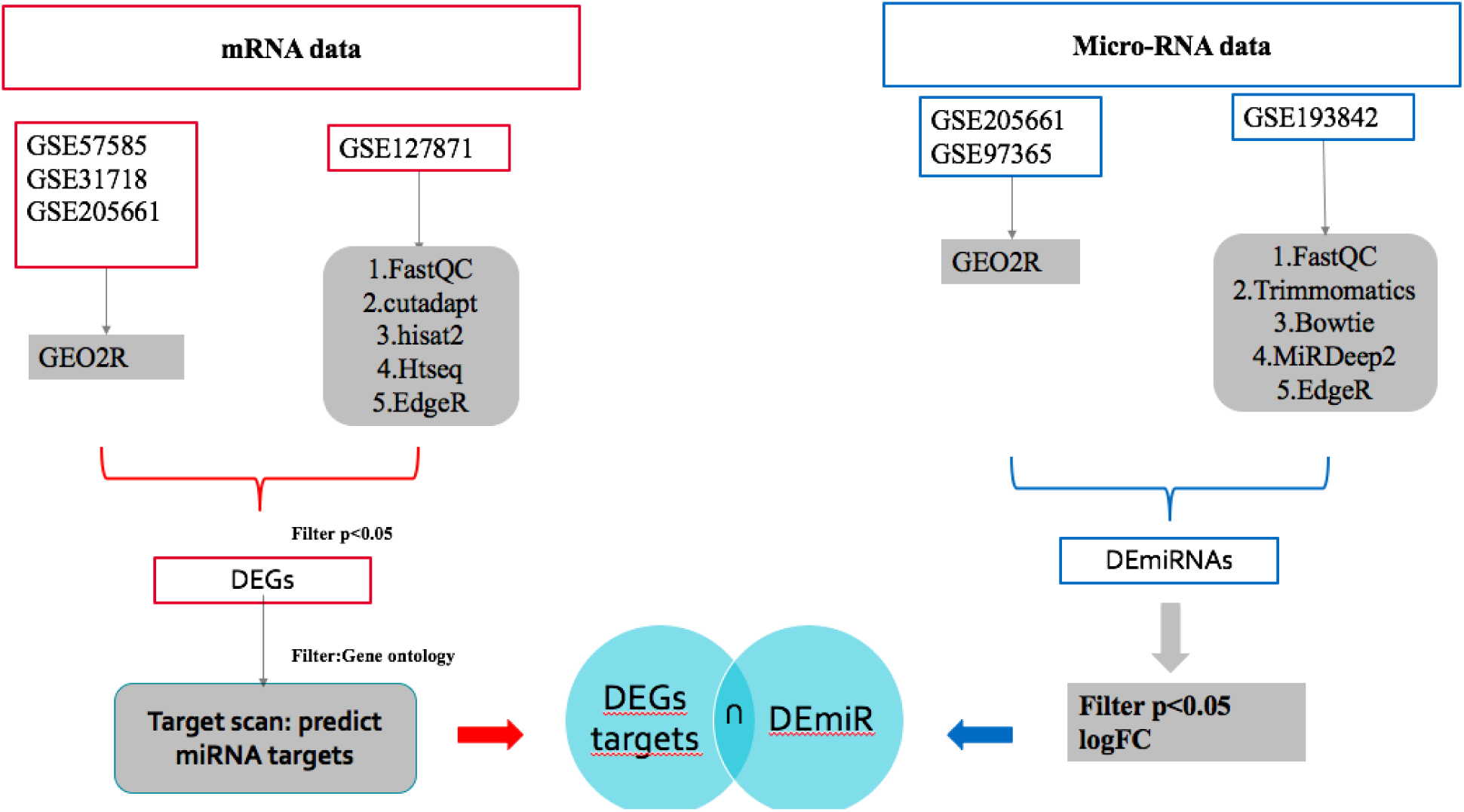
Data analysis workflow of mRNA and miRNA datasets.

#### 2.3 Differential Expression Analysis of mRNA and microRNAs

To investigate differentially expressed genes, we chose two tools based on data types: EdgeR [60] for GSE 127871 and GSE 193842 and the GEO2R tool available in the GEO database for GSE205661, GSE97365, GSE57585 and GSE31718 where the following parameters were used: adjustment of p:Benjamini and Hochberg values (false discovery rate); apply Limma precision: yes; force normalization: yes.

Subsequently, we selected only significant genes (*p-value* lower or equal to 0.05), and then filtered by logFC to obtain overexpressed (logFC>0) and underexpressed (logFC<0) genes.

#### 2.4. Gene Ontology Analysis

According to the gene ontology, we set a list of mRNAs that can be directly related to the disease. By using EnrichR version 3.1, gene ontology analysis and gene set enrichment analysis was carried out [40] to identify important pathways represented in the expression data.

#### 2.5 Prediction of Target Genes

We used the Target scan web tool [76] for the prediction of microRNAs targeting significantly expressed mRNAs. Candidate microRNAs obtained by Target scan are subsequently compared with the results of microRNA analyses in the different datasets to check whether they can regulate the target genes.

## Results

### 1. mRNA analysis

#### 1.1 GSE 57585

In our study, there were two groups of.TLE patients. The first composed of the early-onset patients who developed the disease shortly after initial precipitating insult. The second group includes the people with late-onset illness who were diagnosed beyond the age of 13. Three males and four females made up both groups.

The comparisons of gene expression profiles showed the absence of a notable distinction between the 2 categories of patients. According to the p-value, none of the genes had a p-value <0.05 (Fig. 2).

**Fig. 2.**
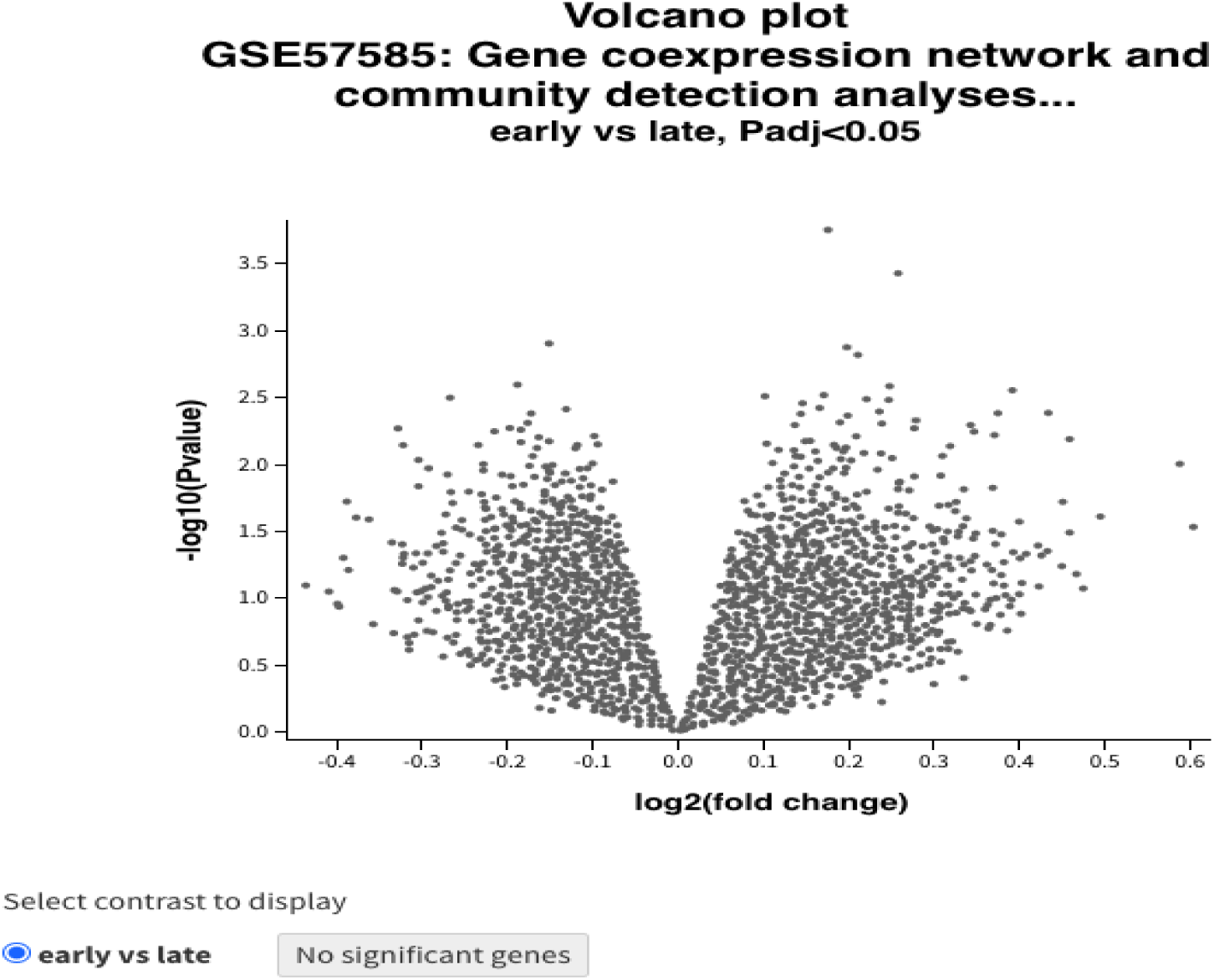
A volcano plot showing DEGs between patients with early and late-onset seizures.

#### 1.2. GSE 31718

This study presents gene expression data from 10 patients with neocortical epilepsy, expression data available for epileptogenic and irritative areas.

In this analysis we are interested in comparing the analysis of epileptogenic zones and irritative zones of brain tissue samples .

We obtained a result of 22,420 genes. Only 61 of them are significant with p value <0.05, of which 46 genes were underexpressed (logFC<0) and 15 genes were overexpressed (logFC>0; Fig. 3).

**Fig. 3.**
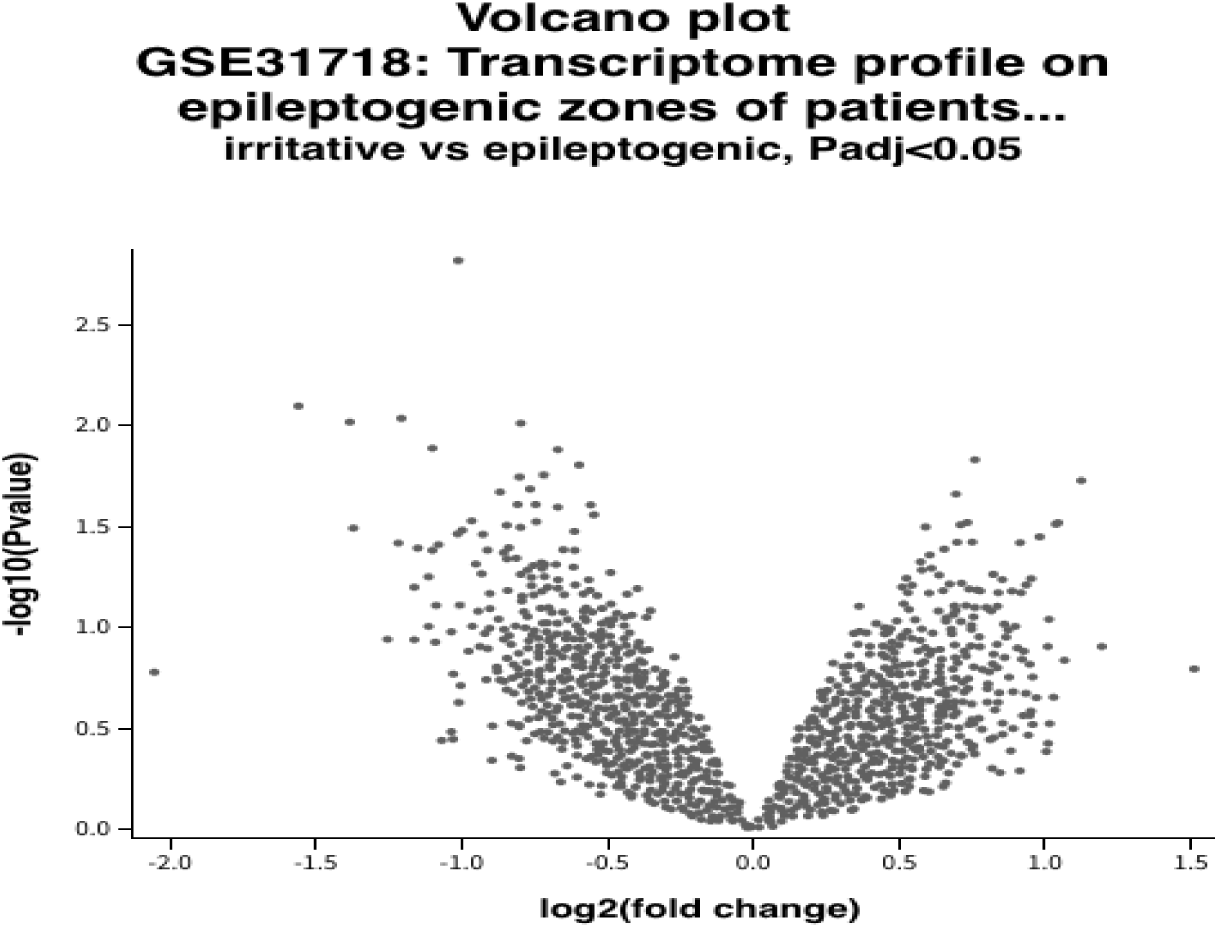
A volcano plot showing DEGs between epileptogenic zone and irritative zone.

These genes were filtered according to the gene ontology in order to find which one can be related to the disease. Any aberrant expression of these particular genes may lead to a trouble of ion transmission which causes an imbalance of charges.

We found that SIX Homeobox 4 (SIX4) and KCTD7 are 2 candidate genes that are underexpressed in the epileptogenic areas.

#### 1.3. GSE 127871

In this study, we investigated expression of mRNA in 12 brain tissue samples of patients with TLE. Patients are divided according to the frequency of occurrence of epileptic seizures into two groups:

- The first group includes patients with a low frequency of epileptic seizures (LFS): SRR8669931, SRR8669932, SRR8669933, SRR8669934, SRR8669935, SRR8669936 (mean =4 seizures/month).
- The second group includes patients with a high frequency of seizures (HFS) : SRR8669937, SRR8669938, SRR8669939, SRR8669940, SRR8669941, SRR8669942 (mean=48 seizures/month).

After analyzing the mRNA expression, we obtained 20 significant genes with p value <0.05 ( supplementary Table 3) among which 16 genes were overexpressed (GBP2, GADD45A, ANKRD22, LIPM, MGP, CASR, MMP3, CALCB, IGKC, TSIX, MUC13, IGHGP, IGHG1, IGHG3, NPS, IGHV1-69D ) and 4 genes underexpressed (KRT17P1, SNORD17, SCARNA10, SEMA3E).

For the overexpressed genes, the logFC values vary between 1.67 (GBP2) and 8.82 (IGHV1-69D) (Fig. 4). For the underexpressed genes, the logFC values vary between -3.06 (KRT17P1 ) and -1.72 (SEMA3E) (Fig. 4).

**Fig. 4.**
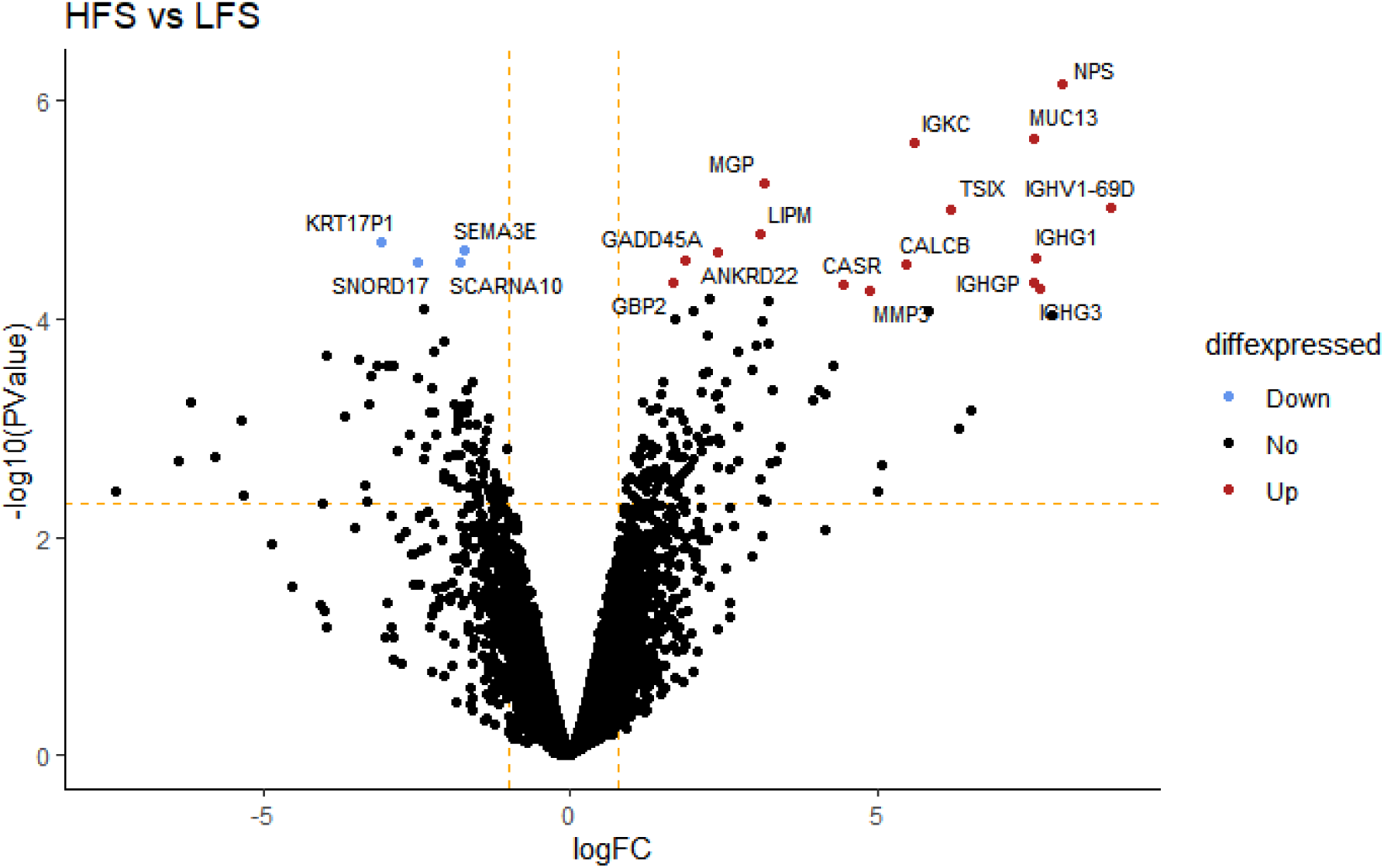
A volcano plot showing DEGs between patients with high frequency of seizures (HFS) versus low frequency of epileptic seizures (LFS). Red dots show over-expressed mRNAs in HFS, black (not significant), and blue shows under-expressed mRNA in HFS.

Functional analysis of significantly expressed genes was done to determine the biological process and pathways linked to the genes identified. This analysis revealed that cell activation, adaptive immune response, immune effector process, activation of immune response were some of the biological processes that are prominently enriched for this set of genes while nucleobase-containing compound metabolic process, RNA metabolic process, cellular nitrogen compound metabolic process were suppressed (Fig. 5).

**Fig. 5.**
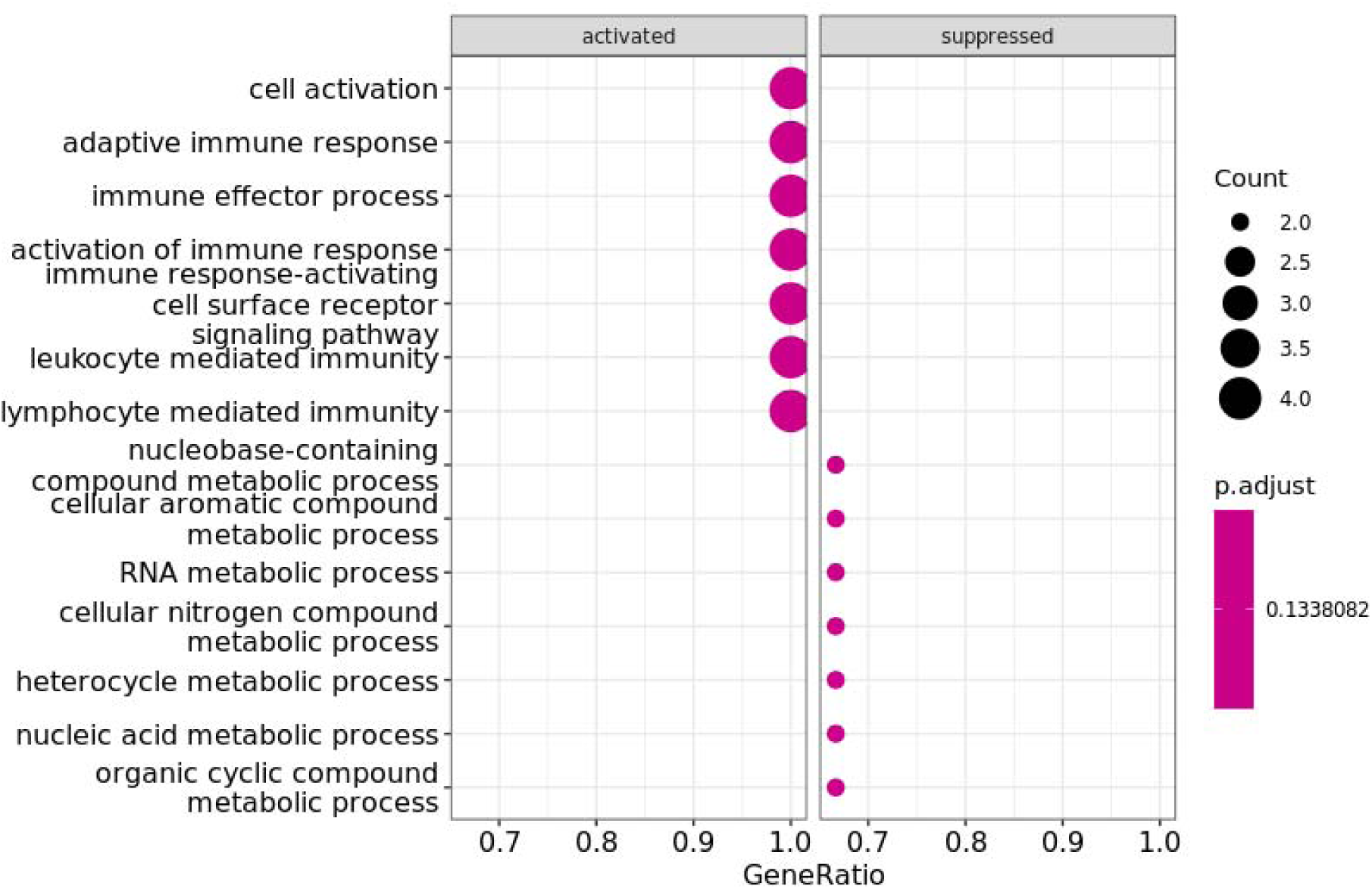
GO biological process for significantly expressed genes.

Neuroactive ligand-receptor interaction pathway was activated while NOD-like receptor signaling pathway and transcriptional misregulation in cancer were observed to be suppressed (Fig. 6A). CALCB and NPS genes which are both upregulated in this study are linked with activated neuroactive ligand-receptor interaction pathway, while GADD45A and GBP2 genes which are also observed to be significantly overexpressed in this study are associated with the suppressed transcriptional misregulation in cancer and NOD-like receptor signaling pathways, respectively (Fig. 6B).

**Fig. 6:**
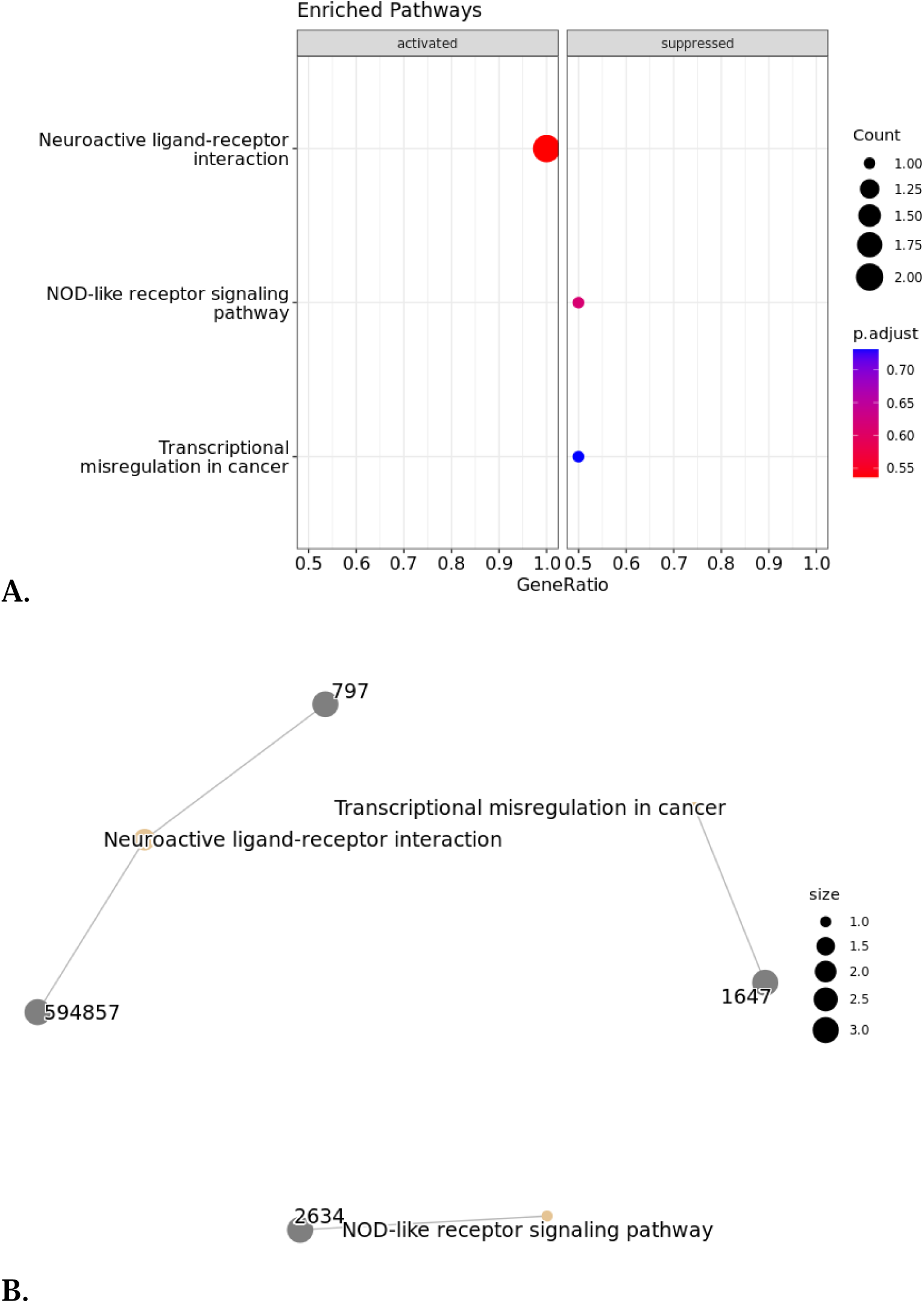
**A:** KEGG pathways for significantly expressed genes. **B:** cnet plot showing the link between genes and the KEGG pathways.

#### 1.4. GSE 205661

In order to explore the difference of mRNA expression in epilepsy, samples were placed in two categories: TLE group (N=6) and control group (N=6).

- Patients: GSM6216213,GSM6216214,GSM6216215,GSM6216216,GSM6216217,GSM6216218.
- Controls: GSM6216219,GSM6216220,GSM6216224,GSM6216225,GSM6216226,GSM6216227.

372 significantly expressed genes were identified, including 271 over-expressed and 101 under-expressed genes.

The logFC varies between 0.111 (H3C3) and 2.589 (B3GAT2) for the upregulated genes, and between -2.908 (MZT2B) and -1.6 (ARHGEF15) for the down-regulated genes.

After filtering the results according to the gene ontology, we observed that NGEF, CABP1, SLC35G1, KCNK3, SLC20A1, SCN1B, SLC2A8, SLC38A7, RIMBP2, CIB3 were down-regulated and SLC4A4, SLC35A5, SLC6A9, NTSR2, KCNN3, S100B, GABARAPL2 were up-regulated.

Subsequently, functional analysis of the selected genes was performed using the EnrichR package to determine the biological processes and pathways associated with the genes. This analysis revealed that organic anion transport, primary metabolic process, organic substance metabolic process, transmembrane anion transport, membrane transport, anion transport and metabolic process were activated by while response to stimulation, regulation of catalytic activity, cellular localization, nervous system process, downregulation of biological processes, downregulation of cellular processes and regulation of molecular functions have been repressed (Fig. 7).

**Fig. 7:**
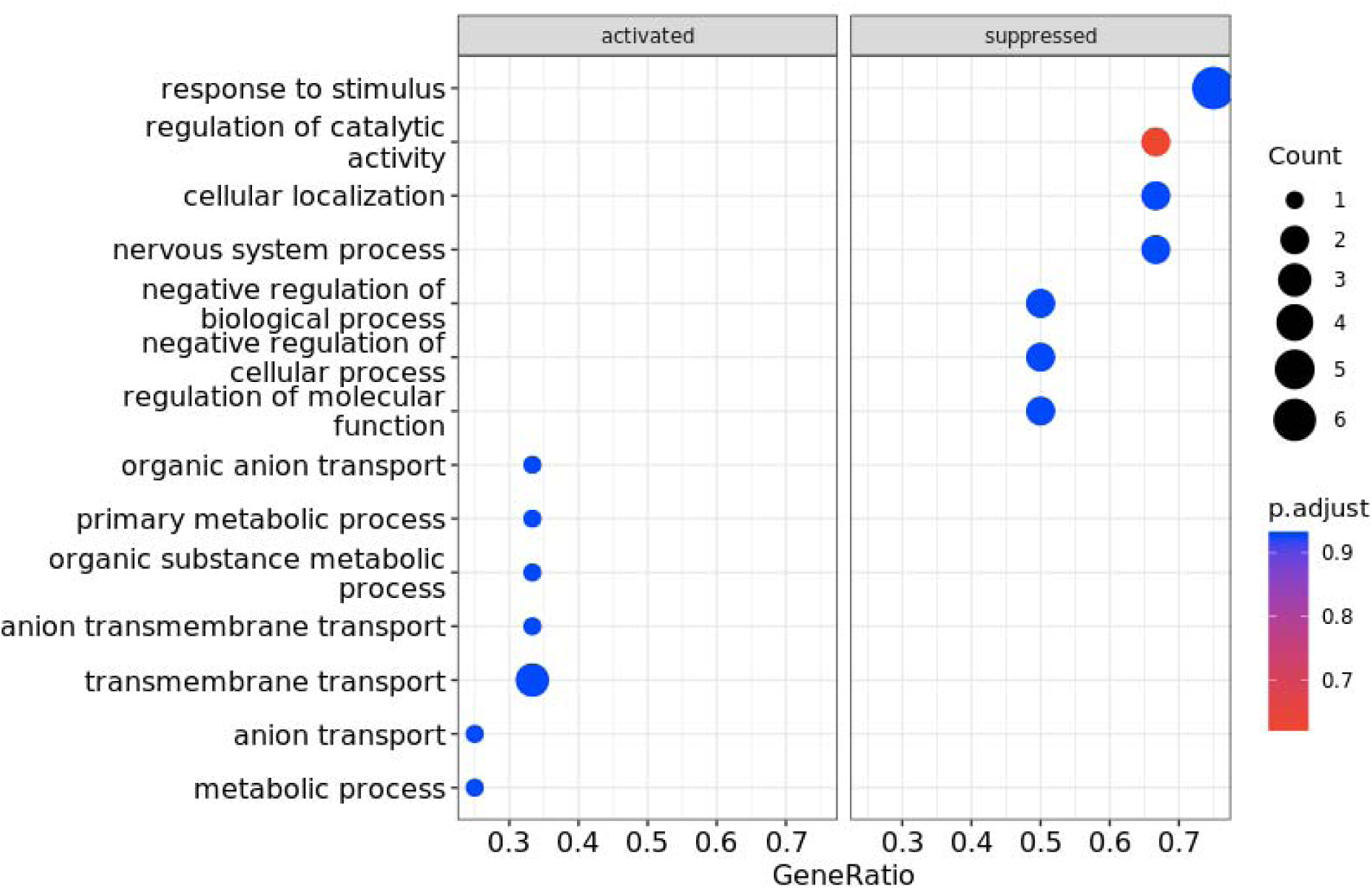
GO biological process for genes detected in the GSE205661 dataset.

These results showed that the proximal tubule bicarbonate salvage, pancreatic secretion, bile, insulin and GnRH secretion pathways, FoxO signaling pathway, autophagy and mitophagy were activated. While the axonal guidance signaling pathway, the aldosterone and cortisol synthesis and secretion pathways and the Cushing syndrome signaling pathway were inhibited (Fig. 8A). The SLC4A4 gene, which was over-expressed in this study, was associated with the activation of the bile secretory pathway, the pancreatic secretory pathway and the proximal tubule bicarbonate scavenging pathway, while the gene KCNN3, which was also significantly over-expressed in this study, was associated with insulin and GnRH secretion (Fig. 8B).

**Fig. 8:**
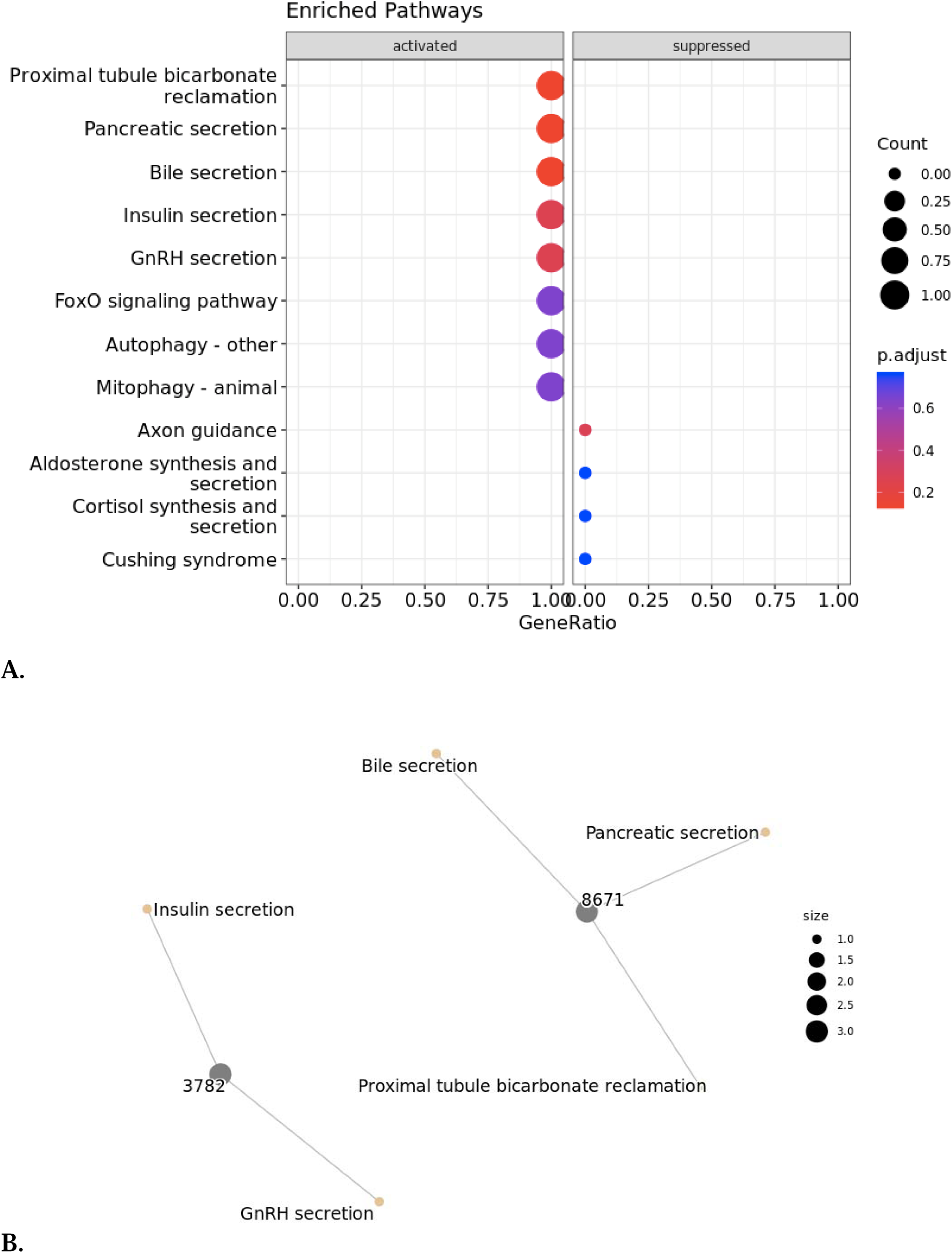
Signaling pathways associated with differentially expressed genes. **A:** KEGG pathways for genes significantly expressed and selected by function in the GSE205661 dataset **B:** Cnet plot showing the link between KEGG genes and pathways in GSE205661.

### 2. microRNA analysis

#### 2.1. GSE 205661

In this study, the comparison of microRNA expression levels is based on 6 patients with TLE with hippocampal sclerosis patients and 6 controls. The choice is based on sex and age in order to have the closest composition to that of the patients.

- Patients: GSM6216213,GSM6216214,GSM6216215,GSM6216216,GSM6216217,GSM6216218.
- Controls: GSM6216219,GSM6216220,GSM6216224,GSM6216225,GSM6216226,GSM6216227.

Results revealed that 34 microRNA were significantly expressed; 19 of them were overexpressed and 15 were underexpressed.

#### 2.2 GSE 97365

This study analyzed expression of microRNA from patients with cortical developmental defects that present a primary cause of central nervous system disorders including epilepsy.

The comparison of microRNA expression levels is based on two groups with the same gender composition: 4 males and one female:

- 5 patients with focal cortical dysplasia type II: GSM2562985, GSM2562988, GSM2562989, GSM2562991, GSM2562992.
- 5 controls: GSM2562979, GSM2562980, GSM2562981, GSM2562982, GSM2562983.

Results revealed that 70 microRNA showed a significant difference with p value <0.05, of which 34 were overexpressed and 36 were underexpressed.

#### 2.3 GSE 193842

In this study, peripheral blood samples were collected from drug-resistant epileptic children, epileptic children who respond to treatment, and healthy children.

Patients were divided into two groups: The first group included healthy controls, who were not affected by the diseases. The second group included epileptic patients (Supplementary Table 1)

After analyzing the miRNA expression, we obtained 80 significant genes with p value <0.05 among which 67 miRNA were overexpressed and 13 miRNA were under-expressed (Supplementary Table 2).

For the highly overexpressed genes, the logFC values vary between 10.60 (hsa-miR-1260a) and 4.50 (hsa-miR-215-5p) (Fig. 10). For the underexpressed genes, the logFC values vary between -2.80 (hsa-miR-548ad-5p) and -1.48 (hsa-miR-5684) (Fig. 9).

**Fig. 9:**
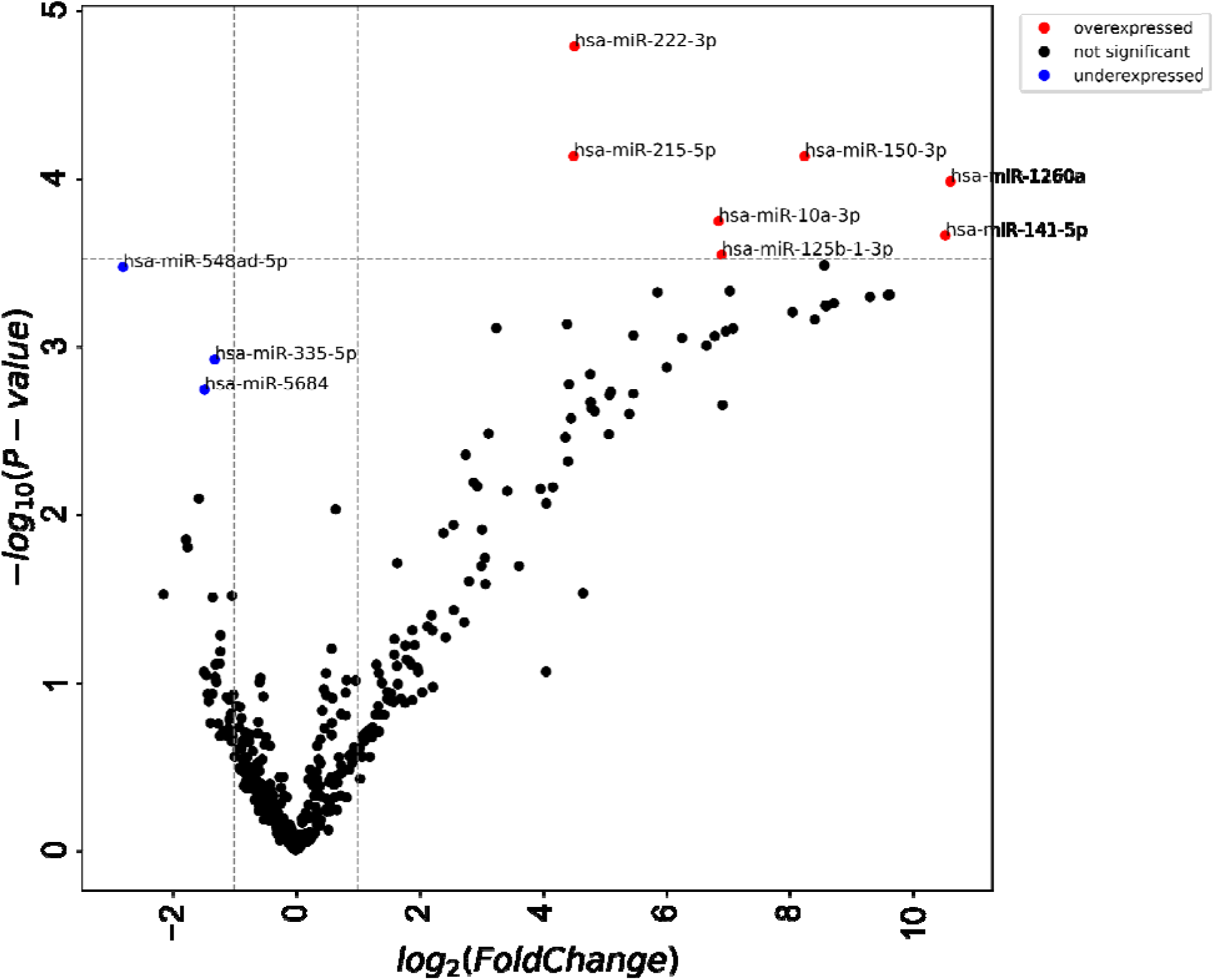
A volcano plot showing DEGs between children with epileptic disease (diseased) versus healthy children (healthy).

**Fig. 10:**
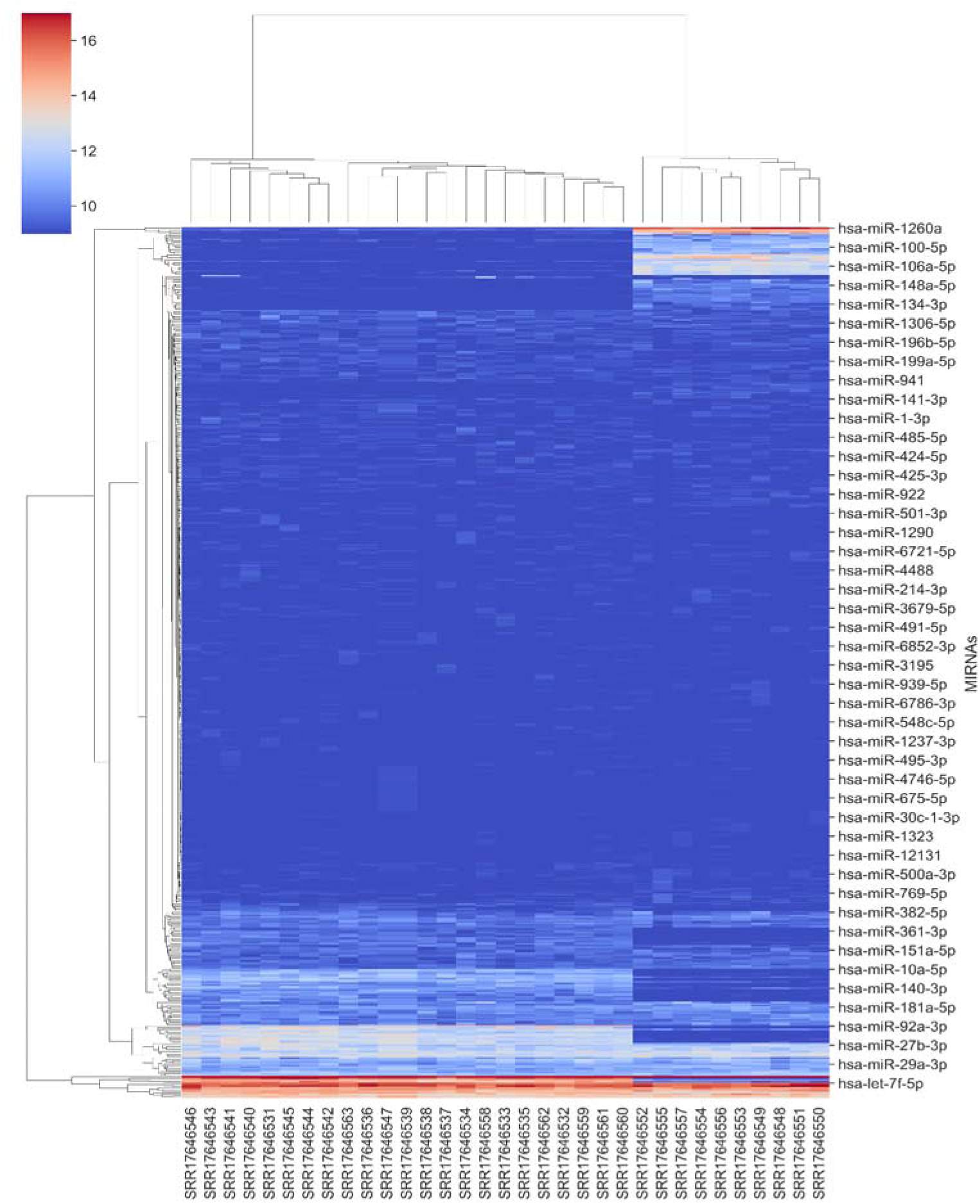
The expression profile of different miRNAs detected (y-axis) against each of the samples (x-axis). The intensity of color changes from blue to red indicating the shift from under expressed miRNAs to over expressed miRNAs, respectively.

Moreover, a heat map was constructed applying the miRNA genes in order to better visualize and understand the changes in miRNA expression between the two groups of children (Fig. 10). Generally, healthy patients and diseased (epileptic) patients clustered distinctly. The expression profile of the diseased samples was different from the healthy patients for 3 genes: hsa-miR-1260a, hsa-miR-100-5p, and hsa-miR-106a-5p: but similar for the other miRNA genes.

Three genes (hsa-miR-1260a, hsa-miR-100-5p, and hsa-miR-106a-5p) are overexpressed for epileptic patients but underexpressed for healthy patients.

## 3. MicroRNA gene target prediction

• GSE127871

For the GSE127871 dataset, the expression of mRNAs between low and high seizure frequency patients was analyzed.

Subsequently, we selected two datasets GSE97365 and GSE193842, to test which of the miRNAs predicted as mRNA targets were differentially expressed at the brain level as well as the plasma level.

The results showed that only two mRNAs were significantly expressed and were targets of miRNAs identified in the GSE97365 dataset. MGP and CASR were overexpressed and predicted as targets of miRNAs that were underexpressed (hsa-miR-155-5p, hsa-miR-195-5p) (Table 2).

**Table 2:** Interaction between miRNAs (GSE97365) and mRNA (GSE127871).

| MiRNAs | Gene target | Gene description |
| --- | --- | --- |
| hsa-miR-155-5p | MGP | Matrix Gla protein |
| hsa-miR-195-5p | CASR | calcium-sensing receptor |

Considering only the differentially expressed miRNAs in GSE193842, 1,334 candidate miRNA-mRNA pairs were predicted using Target scan.

A total of 567 mRNAs were target genes for under-expressed miRNAs. Similarly, 767 mRNAs were target genes for overexpressed microRNA genes. Most of the miRNAs interacted with more than one mRNA gene, indicating their potential to regulate the expression of multiple genes. Interestingly, some mRNAs that had a significant influence on the disease were also predicted as targets for our miRNAs. Three overexpressed mRNAs (GBP, MGP) were predicted as potential targets for three underexpressed miRNAs (hsa-miR-335-5p, hsa-miR-3065-5) (Table 3). Also one mRNA that was under-expressed: SEMA3E was predicted as a potential target for a miRNA that was over-expressed (hsa-miR-1260a) (Table 3).

**Table 3:** Interaction between microRNA (GSE193842) and mRNA (GSE127871).

| Micro-RNAs | Gene Target | Gene description |
| --- | --- | --- |
| hsa-miR-335-5p | GBP2 | Guanylate Binding Protein 2 |
| hsa-miR-3065-5p | MGriPMat | x Gla protein |
| hsa-miR-1260a | SEMA3E | semaphorin 3E |

• GSE31718

For the dataset GSE31718, we investigated differential mRNA expression between brain samples from epileptogenic and irritative areas of patients with drug-resistant neocortical epilepsy.

The GSE97365 dataset was chosen to test which of the miRNAs were regulators for the differentially expressed mRNAs.

The results showed that only one mRNA was differentially expressed and the target of differentially expressed miRNAs in the GSE97365 dataset.

KCTD7 was underexpressed and predicted to be a target of hsa-miR-155-5p that was overexpressed (Table 4).

**Table 4:**
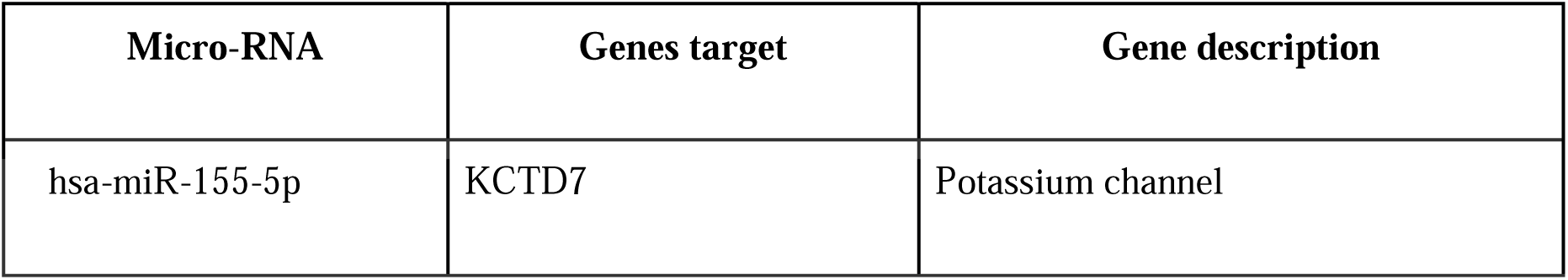
Interaction between miRNA (GSE97365) and mRNA (GSE31718)

Subsequently the mRNA expression data were compared with the results of significantly expressed miRNAs in plasma (GSE193842). The results showed that the miR-759 that was overexpressed in epilepsy patients is a regulator of the SIX4 gene (Table 5).

**Table 5:** Interaction between microRNA(GSE193842) and mRNA(GSE31718)

| Micro-RNA | Genes target | Gene description |
| --- | --- | --- |
| hsa-miR-759 | SIX4 | SIX Homeobox 4 |

• GSE205661

For the data set GSE205661, DGE was investigated for mRNA and miRNA between patients and controls, and the target scan online program was used to predict the mRNA targets.

Results have shown that a total of 5 mRNAs were significantly expressed and were targets of identified miRNAs (Table 6). Three mRNAs (CABP1, SLC20A1, SLC35G1) were downregulated and predicted as targets of upregulated miRNAs. Two mRNAs (KCNN3, SLC4A4) were upregulated and predicted as targets of down regulated miRNAs (Table 6).

**Table 6:** GSE205661 miRNA-mRNA gene targets.

| microRNA | Gene target | Gene description |
| --- | --- | --- |
| hsa-miR-27a-3p | CABP1 | calcium binding protein 1 |
| hsa-let-7b-5p | SLC20A1 | solute carrier family 20 member 1 |
| hsa-miR-15a-5p<br>hsa-miR-195-5p | SLC35G1 | solute carrier family 35 member G1 |
| hsa-miR-149-5p<br>hsa-miR-132-3p | KCNN3 | potassium calcium-activated channel subfamily N member 3 |
| hsa-miR-149-5p | SLC4A4 | solute carrier family 4 member 4 |

After comparing the miRNA results in all the data sets, we found that hsa-miR-1260a is upregulated in GSE205661 and GSE193842 (2 datasets comparing patients and controls in two different tissues), hsa-miR-155-3p is upregulated in GSE 193842 and GSE97365 and hsa-miR-149-5p is down-regulated in GSE205661 and GSE193842.

## Discussion

TLE is a subtype of epilepsy that may spread through a network of neuronal connections to the adjacent brain tissue [57]. The age of epilepsy onset is a significant part in the classification of idiopathic epilepsy, and a crucial factor in the determination of the disorder subtype leading to gene identification [36].

In the gene expression analysis of early onset compared to late onset patients (onset later than thirteen years). No genes were significantly differentially expressed. A recent study revealed distinct molecular pathways of each group of patients [36]. <u>Moreira-Filho *et al*</u>., [51] reported that the early onset epilepsy severely touches the ictal hippocampus but it’s associated with a mechanism that allows the compensation. In late onset epilepsy, the brain is no longer plastic and becomes less able to adopt the compensatory mechanism [51]. Despite the lack of clear distinct differences in the genetic makeup of the two conditions, it has been suggested that early therapy as well as infant seizure control be implemented as soon as possible, as uncontrolled seizures have been shown to impede brain function, with the consequences being more acute in children and less severe as people get older when the seizure first starts [25]. In a study by Hsu [34], young children with image-confirmed arterial ischemic stroke where the cortex was involved were more prone to experience early-onset seizures [34]. Interestingly, after the acute episodes, 65 % of kids with early-onset seizures also experienced late-onset seizures [34].

Gene expression data comparing expression between epileptogenic zones and irritative zones of ten people with drug-resistant neocortical epilepsy reveals that SIX4 and KCTD7 were significantly underexpressed. These genes are important because SIX4, based on its particular expression in specific neural cells and skeletal muscles; it is assumed to be important in neurogenesis [55]. This implies that the underexpression of SIX4 could lead to impairment in neural development [55]. Through vertebrate evolution, the SIX4 expression pattern has generally remained constant [55]. Similarly, KCTD7 was linked with progressive myoclonic epilepsy in a study that quantified histological images of different mouse strains [23]. Severe epilepsy has been associated with KCTD7 gene mutations and regression of cognitive functions in infants due to alterations in bipolar neurons [77], [23]. Bipolar cell integration into developing synapses may be impacted by Kctd7-dependent variations in neuronal activity [23]. Underexpression of these genes in our study population proves their role in the disease and thus could be potential biomarkers for treatment development for people with neocortical epilepsy and are drug-resistant.

Gene expression analysis of TLE patients (GSE127871) who experienced high frequency seizures (mean = 48 seizures per month) compared to low frequency seizures (mean = 4 seizures per month) provides evidence that neuroactive ligand-receptor interaction pathway (activated), NOD-like receptor signaling pathway and transcriptional misregulation in cancer pathway which were suppressed are involved in temporal lobe epilepsy. By attaching to intracellular receptors, which have the potential to bind transcription factors and control gene expression, neuroactive ligands have an impact on how neurons operate [70]. Memory performance is diminished when the genes involved in neuroactive ligand-receptor interaction are disrupted [70]. Moreover, according to Liu *et al.,* [45] unnatural interactions between neuroactive ligands and receptors increase the susceptibility to epileptic seizures. Therefore, our findings show that CALCB and NPS genes were over-expressed and were predominately enriched in the neuroactive ligand-receptor interaction pathway, resulting in neuro dysfunction. This pathway and the genes associated with this pathway could be a novel focus for examining the mechanisms of temporal lobe epilepsy.

The study of expression of TLE patients vs normal highlights a unique set of differentially expressed genes and enriched KEGG pathways. FoxO signaling pathway is activated for these sets of genes; Kim *et al.,* [39] discovered a connection between seizure-induced neuronal death among temporal lobe epilepsy patients and Akt-FoxO3a signaling [39]. These findings closely match our findings of the KEGG analysis.

In another study, we focused on comparing mRNA (GSE205661) expression profiles between HS-TLE patients and controls. Our analysis allowed us to highlight the involvement of the activated pathways : proximal tubule bicarbonate reclamation , pancreatic secretions, bile, insulin and GnRH secretion, the FoxO signaling pathway, autophagy and mitophagy in temporal lobe epilepsy associated with hippocampal sclerosis. Menzies *et al.* [75] reported that autophagy dysfunction plays a role in the development of neurodegenerative diseases such as Alzheimer’s or Parkinson’s disease. Additionally, previous studies have proven that altered mitophagy contributes to synaptic dysfunction and cognitive impairment [74].

The epigenetic regulation of genes is attributed to more than one process. In this work, we are interested in microRNAs as regulators of gene expression. Viewing the fine balance between mRNA and miRNA in the cell, any abnormal expression of a specific miRNA can affect several genes and pathways, leading to an imbalance in the cell function and resulting in pathologies. That is why we performed combinatorial analysis where we used a prediction software named Target Scan in order to reveal the targets of a list of altered genes in epilepsy.

In this research, miR-27a-3p was upregulated in TLE patients compared to the controls. This miRNA was predicted as a target for calcium binding protein 1 (CABP1) that was down-regulated. A previous study had confirmed that miR-27a-3p is associated with epilepsy and it is suggested to be a potential biomarker for TLE [58]. Another study has shown that the inhibition of miR-27a-3p leads to the prevention of epilepsy-induced inflammatory responses and the neuronal apoptosis by the inhibition of MAP2K4 expression [46]. Loriatti *et al.,* [47], reported that the expression level of miR-27a-3p was elevated in serum but it failed to differentiate mTLE patients from controls.

MiRNA expression analysis of TLE patients (GSE205661) revealed that hsa-miR-15a-5p and hsa-miR-195-5p were upregulated and targeting a downregulated gene: solute carrier family 35 member G1 (SLC35G1) whereas hsa-miR-15a-5p was downregulated in epilepsy patients compared to controls in other studies.

A previous study reported that has-miR-15a-5p was under-expressed in epilepsy patients with diagnostic values of 80% sensitivity and 73% specificity. Their results showed that this microRNA plays a role in the development of epilepsy by regulating the inflammation or apoptosis [15]. Consistently, the downregulation of miR-15a-5p expression levels in serum have been reported in children with TLE by [44]. They also investigated the effect of the over-expression of this microRNA on the cell viability and apoptosis of hippocampal cells and their results proved that high levels of miR-15a-5p caused the inhibition of apoptosis and enhanced the cell viability. These results bring out the important role of miR-15a-5p in the progress of TLE.

A previous study evaluated the whole human hippocampal miRNome in order to identify microRNAs implicated in the development of mTLE associated with hippocampal sclerosis [26]. They found that miR-195-5p was upregulated which supported our results and other previous research that evaluated microRNA expression in HS-TLE patients [72].

The Solute Carrier family is a family of genes coding for membrane transporters in the endothelial cell; they play a crucial role in maintaining the concentration of the brain’s solutes [66]. They are highly expressed in the brain and they are important in the energy production of normal neurons. The Solute Carrier family was associated with many neurodegenerative disorders such as schizophrenia, epilepsy, anxiety and depression [24].

In the current study, SLC35G1 and SLC20A1 were downregulated in epileptic patients compared with controls .SLC35G1 was a target of the two upregulated miRNAs: miR-15a-5p and miR-195-5p and the gene SLC20A1 was a predicted target of hsa-let-7b-5p which was upregulated. Previous studies have suggested that has-let-7b-5p was associated with multiple cancers [73], [61]. It promotes cell apoptosis in Parkinson’s disease by targeting HMGA2 which is responsible for the mediation of gene expression, but no previous evidence supports the association of has-let-7b-5p with epilepsy [29], [35]. These results demonstrate the importance of this gene family in the mechanism of the disease and the implication of these miRNAs in the progress of epilepsy.

In this study, hsa-miR-149-5p was downregulated in patients compared to control for both brain and blood tissues at the same time (GSE205661 and GSE193842 datasets). It is supposed to regulate the SLC4A4 and KCNN3 which were upregulated in patients.

A previous study reported that the downregulation of hsa-miR-149-5p was found during a brain ischemia in rats and the upregulation of this miRNA regulates the caspase3 implicated in the mediation of apoptotic neuronal cell death via Sirt1/p53 axis [67].

The downregulation of miR-149-5p affects the expression levels of KCNN3 in hippocampal cells, and our results showed that KCNN3 was upregulated in patients of GSE205661 but previous studies reported that the expression level of KCNN3 was decreased in pilocarpine-treated epileptic rats and that the use of a gene blocker increases the excitability [53]. This evidence supports the role of KCNN3 in controlling neuronal excitability and suggests that the loss of function of these potassium channels may lead to increased epileptic seizures. We also found that KCNN3 was a target for miR-132-3p that was downregulated in TLE patients, so we can conclude that the upregulation of KCNN3 results from the downregulation of its two regulators miR-149-5p and 132-3p.

The aberrant expression of miR-132-3p was reported in many other neurodegenerative diseases. For instance, Johnson *et al.,* [37] reported that miR-132-3p was downregulated in Huntington’s model mice compared with controls and that the downregulation leads to increases in its target expression. Also, Miller *et al.,* [50] reported that miR-132-3p was downregulated in brain samples of patients with schizophrenia while the predicted gene targets were upregulated.

Our results showed that miR-155-3p was differentially expressed in hippocampal tissue and blood samples. The analysis of GSE205661 and GSE193842 indicated that miR-155-3p was significantly higher in TLE patients compared to controls but none of its gene targets predicted by Target Scan was downregulated.

miR-155 has been shown to be upregulated in epileptic patients in various studies [30], [49]. It showed significant biomarker performance with 73% sensitivity and 80% specificity [13]. This evidence is consistent with our results but we suggest that miR-155-3p may be regulating downregulated genes in GSE205661 and GSE193842 that are undiscovered yet.

In this research (GSE193842), miR-335-5p was downregulated in children suffering from epilepsy compared to controls. This microRNA was predicted to target the mRNA gene guanylate binding protein 1 (GBP) that was up-regulated in patients with TLE with hippocampal sclerosis. Previous studies have shown similar expression profiles of the miR-335-5p genes in epilepsy disease [52], [63], [69]. Hence, the potential of this miRNA gene is not yet clearly understood in relation towards epileptic diseases. However it has been linked to the neurological disorder such as Alzheimer’s disease through the inhibition of β-Amyloid (Aβ) by targeting c-jun-N-terminal Kinase [71]. GBP is linked to inflammation through the inhibition of endothelial cell proliferation and by protecting the cells from interferonγ (IFNγ)-induced apoptosis [33]. Studies [43] have experimentally confirmed the potential of miR-335-5p in regulating GBP expression profiles.

The MiR-942-5p was down regulated in children suffering from epileptic diseases compared to controls. This miRNA was predicted to target the mRNA gene, a calcium sensing receptor (CASR) that was up-regulated in patients with TLE with hippocampal sclerosis. The CASR gene is expressed in the central nervous system and the role it plays in epilepsy has been postulated in previous studies. High levels of the CASR gene have been associated with the disease [59]. It has been hypothesized that the epilepsy could be caused by alteration in serum [Ca^2+^] due to disruption in the parathyroid hormone (PTH) level, which leads to suppressing inhibitory neurons, activating excitatory neurons, or a more general effect on certain brain circuitry [62].

Intriguingly, this mRNA gene has been shown to have a significant expression profile in the GSE127871 study samples. Moreover, the results of our computational analysis indicated that the miR-942-5p, a down-regulated miRNA gene, was a non-conserved seed region target for the CASR gene. Hence, we postulate that the miR-942-5p gene could play a potential role in regulating this gene. However, other studies involved in cancer cell lines have shown other miRNAs such as miR-21, miR-145, and miR-135a, to have a negative correlation to the expression of CASR [31], [64], [65].

In our review, miR-342-5p was upregulated in children with epilepsy compared to the healthy children. This was replicated in a study by Wang *et al.* [68], where miR-342-5p was among the hypo-expressed genes among people with treatment-resistant epilepsy versus patients who respond well to anti-epileptic medicines and control subjects. Hence, we also exploited the potential target of this miRNA and identified ankyrin repeat domain 22 (ANKRD22) as a potential candidate, because of the inversely correlated expression between the gene and the miRNA and the interaction between the two genes. Interestingly, the GSE127871 study postulated a low expression profile of the mRNA gene by comparing patients with TLE with the hippocampal patients. MiR-1260a was shown to have high expression profiles in GSE193842. Conversely, this phenomenon was observed in the neocortex of patients with mesial temporal lobe epilepsy compared to control [54] In this study, we hypothesized that miR-1260a, an up regulated miRNA gene was a non-conserved seed region targeting the Sema3E gene.

## Conclusions and next steps

This research was designed *in-silico* to study the transcriptomic profiles of different epilepsy patients. We identified multiple differentially expressed genes that we suggest are a part of the pathophysiology and development of epilepsy. Additionally, we identified miRNAs which are potential biomarkers that are regulating the expression of mRNAs involved in epilepsy. MiR-155-3p was upregulated in our study, we postulate that it could be targeting a downregulated gene that is yet to be discovered. The function of the miRNA gene and the target protein coding gene is also yet to be discovered. The following miRNA genes: MiR-335-5p, miR-942-5p ,and miR-1260a were identified in this study as regulators of known epilepsy-associated genes BP , CASR, and Sema3E, respectively. Some of our identified biomarkers have already been validated in other studies but *in-vitro* studies may be needed to corroborate some of our findings. Also, the data analyzed correspond to neuronal cells of hippocampal tissue and Serum Small Extracellular Vesicles and so, studies performed on other cell types might provide other results. Finally, our bioinformatics pipeline could be applied to study gene expression and for the analysis of miRNA data generated for other diseases.

## Declarations

### Availability and Requirements

Project name: e.g. Transcriptomic Profiling of Epilepsy Patients Using Bioinformatics Analysis

Project home page: https://github.com/omicscodeathon/epilepsy_rna

Operating system(s): Platform independent if using the associated Docker/Singularity images.

Mac OS X or Linux CLI if run outside Docker/Singularity images.

Programming language: Bash, R 3.5, Perl, Python.

Other requirements: Python 3.9 or higher, Java JDK, Chrome, Firefox, and Safari web browser.

License: MIT

Any restrictions to use by non-academics: None.

### Availability of data and materials

The dataset analyzed during the current study as a case study is publicly available at https://github.com/omicscodeathon/epilepsy_rna/data. The data supporting the results reported in this manuscript is included within the article and its additional files. The generated progress reports are in HTML format and can be viewed using any preferred browser such as Chrome, Safari, Internet Explorer and Firefox. The Project repository which also includes the entire code and other requirements can be downloaded from https://github.com/omicscodeathon/epilepsy_rna. The guidelines for implementing this tool and related updates, are available at: https://github.com/omicscodeathon/epilepsy_rna/blob/master/README.md.

## Abbreviations

AEDs: Anti-Epileptic Drugs
AMPA: α-amino-3-hydroxy-5-methyl-4-isoxazole propionic acid
ANKRD22: ankyrin repeat domain 22
Aβ: β-Amyloid
CASR: calcium sensing receptor
CLI: Command-Line Interface
DEG: differentially expressed genes
EEG: electroencephalogram
FDR: False Discovery Rate
GABA: gamma-aminobutyric acid
GEO: gene expression omnibus
GUI: Graphical User Interface
HFS: high frequency of seizure
HS-TLE: hippocampal sclerosis with temporal lobe epilepsy
KCTD7: Potassium Channel Tetramerization Domain Containing 7
mRNA: messenger RNA
miRNA: microRNA
mTLE: mesial temporal lobe epilepsy
NMDA: N-methyl-D-aspartate
LFS: Low Frequency Seizure
TLE: Temporal Lobe Epilepsy
SIX4: SIX Homeobox 4
SRA: Sequence Read Archive
PTH: Parathyroid Hormone

## Acknowledgements

The authors thank the National Institutes of Health (NIH) Office of Data Science Strategy (ODSS), the National Center for Biotechnology Information (NCBI) and the Genetics Society of America for their immense support before and during the October 2022 Omics codeathon organized in collaboration with the African Society for Bioinformatics and Computational Biology (ASBCB). The authors thank Brian Musyoka for his assistance in the bioinformatics analysis of the miRNA data and in the drafting of the manuscript.

## Author Contributions

GB and FEA conceived the original idea. MN developed the pipeline. MN performed the bioinformatic analysis of mRNA data, CN performed the bioinformatic analysis of microRNA data. FEA performed the GEO2R analysis. MN, FEA, GB and OIA drafted the manuscript. OIA reviewed the manuscript and supervised the bioinformatics analysis and provided critical feedback that helped improve the work. OIA’s role was also in the administration of the project and provision of the resources to facilitate and complete the analysis. OIA provided guidance, editing and final review of the manuscript. All authors read and approved the final manuscript.

## Declarations

### Ethics approval and consent to participate

Not applicable.

### Consent for publication

Not applicable.

